# Microbiome composition and function vary with depth in the Mediterranean gorgonian *Eunicella singularis*

**DOI:** 10.1101/2025.10.17.683177

**Authors:** Mohammad Binsarhan, Lorenzo Bramanti, Béatrice Rivière, Lidia Montiel, Francisco Latorre, Javier del Campo, Pierre E. Galand, Ramiro Logares

**Author notes:** Correspondence: Mohammad Binsarhan^1,2^ and Ramiro Logares^1, 1^Institute of Marine Sciences (ICM), CSIC, Barcelona, Spain, ^2^ Marine Biology Department, Faculty of Marine Science, King Abdul-Aziz University, Jeddah, Saudi Arabia.

## Abstract

**Background:** The Gorgonian coral, *Eunicella singularis*, is one of the main components of Mediterranean marine animal forests, whose canopies play a key role in Mediterranean sublittoral ecosystems due to their capacity to provide shelter, food, and nursery ground to several species. Like other gorgonian species, *E. singularis* faces environmental threats with potential repercussions on the associated biodiversity. This photophilic octocoral, spanning the Western Mediterranean, Adriatic, and Aegean Seas at depths of 10-70m, engages in symbiosis with *Symbiodiniaceae* dinoflagellates, thereby influencing their resilience in nutrient-poor habitats. However, mesophotic populations (∼60m depth) are characterized by very low *Symbiodiniaceae* density, prompting questions regarding metabolic adaptations. The associations of corals with specific bacteria that differ from those in the surrounding seawater suggest a role in host health and physiology. In particular, in mesophotic colonies, bacteria may perform activities that Symbiodinium typically carries out in shallow colonies. Here, we applied metagenomics techniques to analyze the changes in the microbiome of *Eunicella singularis* with depth, analyzing DNA samples from shallow (12m) and mesophotic (57m) colonies in the Northwestern Mediterranean Sea.

**Results:** High-coverage metagenomes (ca. 80 Gb per sample) were generated from *E. singularis* colony samples. We observed significant changes in the relative abundance of prokaryotic and microbial eukaryotic symbionts with depth. Mesophotic samples exhibited higher levels of symbiotic prokaryotes, dominated by *Endozoicomonas* and *Bermanella*, whereas shallow samples were enriched with the symbiotic dinoflagellate *Symbiodiniaceae*. The metabolic potential of the microbiome also varied with depth. The shallow microbiome showed a prevalence of photosynthesis and carbon fixation pathways. In turn, the mesophotic microbiome exhibited a higher abundance of metabolic functions related to vitamin biosynthesis, energy metabolism, especially carbon metabolism, as well as pathways associated with carbohydrates, amino acids, and cofactors.

**Conclusions:** Our results indicate that the structure and metabolic function of the microbiome of *Eunicella singularis* change with depth. In the absence of symbiotic dinoflagellates, associated bacteria seem to use different sources of carbon, in addition to cycling nutrients and vitamins, which could influence coral health. These findings gain significance in the context of global change, as shifts in oceanic conditions may affect the coral microbiome.

## Introduction

In the Mediterranean Sea, gorgonian corals are the most common engineering species [1] with a key role in sublittoral ecosystems[2]. When their populations are dense enough, they form true forests [3] whose canopies provide shelter, food, and nursery ground for a variety of species, resulting in higher biodiversity associated with dense forests [4]. *Eunicella singularis* (the white gorgonian) is a photophilic octocoral that typically colonizes horizontal to sub-horizontal substrates between ∼10–70 m in the western Mediterranean, Adriatic, and Aegean seas [5,6]. Notably, *E. singularis* is unique as the sole Mediterranean gorgonian species known to harbor symbiotic *Symbiodinium* dinoflagellates [7,8], specifically of the genus *Philozoon* [9]. This symbiotic relationship supplies the coral with energy through photosynthesis-driven metabolites and contributes to the absorption of dissolved nutrients, enabling coral growth also in nutrient-poor environments [10]. On the other hand, the coral provides *Symbiodinium* with phosphorus, inorganic nitrogen, inorganic carbon, and habitat [11].

Depth, however, has a strong influence on this association. Shallow colonies (up to approximately 10m depth) of *E. singularis* can establish a symbiotic relationship with *Symbiodiniaceae*. However, in deeper colonies, the population density of *Symbiodinium* decreases [12], observing a complete absence of it in some cases [13]. This gradient points to a depth-dependent transition from autotrophy to heterotrophy [7], which raises the question of how associated microbial communities contribute to coral nutrition and resilience across depth habitats [8].

Beyond algal symbionts, such depth-related reliance on heterotrophy underscores the potential importance of bacterial partners in sustaining gorgonian metabolism. Yet, our understanding of how microbes contribute to the physiology of *Eunicella singularis*, particularly in the absence of *Symbiodiniaceae*, remains limited [14]. Like other corals, octocorals live in symbiosis with complex microbial communities that contribute to nutrient cycling, pathogen defense, and holobiont fitness [15]. These associated bacteria are distinct from those in the surrounding seawater [15–17]. Investigations of different gorgonian species have highlighted species-specific core bacteria (*i.e.,* bacteria that tend to be present in the microbiome of all coral colonies) [18]. Indeed, *Eunicella* spp. has shown signs of phylosymbiosis with Sphingomonadales [19] and Poribacteria [20].

Depth can be an important factor shaping gorgonian microbiomes, influencing both community diversity and stability. At intermediate depths (∼35 m), *Eunicella singularis* harbors a core microbiome dominated by *Endozoicomonas*, probably acquired horizontally during early developmental stages [21]. Similarly, investigations of *Eunicella cavolinii* across depths of 24, 30, and 41m showed consistent dominance of *Endozoicomonas*, but also revealed depth-related shifts in taxa such as *Aquimarina* and *Myxococcales* [22]. Another study on *Eunicella verrucosa* investigated the impact of depth within a range of 6–27 m [23]. Denaturing gradient gel electrophoresis (DGGE) and clone libraries revealed the prevalence of Gammaproteobacteria, particularly from the genus *Endozoicomonas*, across various depths. The shallow samples (6-9m) displayed a bacterial community that was less similar and more diverse than that of the mesophotic samples [23].

The physiology and fitness of the coral host are closely tied to the diversity and metabolic functions of its microbiome, suggesting that depth-driven changes in microbial composition may directly impact coral performance. However, most research on octocorals has emphasized taxonomic rather than functional variation[20,24]. Functional changes linked to depth-related shifts in coral microbiome community structure remain poorly understood.

Here, we investigate how the diversity and function of *Eunicella singularis’* microbiome changes with depth. We analyzed microbiomes from both shallow (12 m) and mesophotic (57 m) habitats using deep shotgun sequencing. We sought to answer the following questions: (i) Does the composition of dominant microbial species vary with depth? (ii) Are there significant functional differences in the microbiomes across depths? (iii) What are the functional roles of the main members of the microbiome at different depths?

## Material and methods

### Sample collection, DNA preparation, and sequencing

Samples were collected from the Banyuls-sur-Mer coast, South of France, Mediterranean Sea, by scuba diving at different depths (shallow: 12 m, mesophotic: 57 m), in December 2020. Branches (10-15 cm each) of 10 *Eunicella singularis* colonies were sampled (five shallow, E_12m_1 to E_12m_5, and five mesophotic, E_57m_1 to E_57m_5). DNA extraction was performed using the Zymo DNA kit following the standard protocol. The extracted DNA was shotgun sequenced using *Illumina*, aiming for a high coverage per sample (Novaseq 2 × 150 bp, > 80 Gb/sample). Sequencing was done at Macrogen in South Korea. The high sequencing depth aims to obtain more information about the microbiome, as we expect that most reads will belong to the coral. Sequences were deposited in the European Nucleotide Archive (ENA; https://www.ebi.ac.uk/ena) under the accession number PRJEB61841.

### Bioinformatic analyses

Sequence quality checks and adapter removal were performed using Cutadapt [v1.16] (minimum length =75, quality cutoff = 20, nextseq-trim = 20, max-n= 0) [25]. The taxonomic diversity in metagenomes was determined using mTAGs, a bioinformatic tool that performs taxonomic annotation of metagenomic reads that belong to the ribosomal RNA, using a 97% similarity threshold to assign reads to Operational Taxonomic Units (OTUs) [26]. Different databases have been used to assign taxonomy, such as SILVA [SILVA-128] for prokaryotic and eukaryotic reads [27] and the PR2 (Protist Ribosomal Reference) database [4.14.0] [28] for protist reads. In addition, to increase the accuracy of protist assignments [29], we extracted the V4 region from mTAGs and aligned the reads against a specifically curated in-house eukaryotic V4 database [30]. SeqKit [v0.16.0] was used to extract sequences [31], and the eukaryotic mTAGs were then aligned against the PR2 database using USEARCH [v.9.2.64] [32] (id = 0.97, mincols = 105, strand = both, -strand both, -top_hits_only -maxaccepts = 0, maxrejects = 0).

Clean metagenomic reads were assembled using MEGAHIT [v1.2.8] [33] with the *meta-large* preset. Prokaryotic and eukaryotic contigs were identified with EukRep [v0.6.5] [34] using a minimum contig size of 2,000 bp. Eukaryotic exons (hereafter, “genes”) were predicted on eukaryotic contigs with MetaEuk [v1-ea903e5] [35] using the following parameters: metaeuk-val= 0.0001, metaeuk-tcov= 0.6, maximal E-Value= 100, min-length= 40, min-exon-aa= 20. In turn, prokaryotic Open Reading Frames (ORFs; hereafter “genes”) with MetaGeneMark [v3.38] [36] and Prodigal [v2.6.3] [37]. MetaGeneMark predicted both partial and complete genes, while Prodigal predicted only complete genes. Only genes ≥ 250 bp were included in the downstream analyses. Prokaryotic and eukaryotic genes from the different samples were pooled separately, and redundancy was removed by clustering them at 95% identity and 80% alignment coverage of the shorter fragment using the ‘linclust’ [38] algorithm from MMseqs2 [v.9-d36de] [39]. The generated prokaryotic catalog had 731,719 dereplicated predicted genes, while the eukaryotic catalog had 558,223 dereplicated genes, including 151,264 *Symbiodiniaceae* genes. The taxonomic annotation of prokaryotic genes was performed with the taxonomy module from MMseqs2 using the Genome Taxonomy DataBase (GTDB) [v89] [40]. Similarly, the eukaryotic genes were taxonomically classified with the taxonomy module from MMseqs2 using EukProt [v2], a database of genome-scale predicted proteins from various eukaryotes, including Cnidaria [41]. Predicted genes were functionally annotated using Diamond [v0.9.22] [42], with a maximum expected value of 0.1, against the KEGG (Kyoto Encyclopedia of Genes and Genomes) database [v2019-02-11] [43]. Diting [v0.9] [44] was used to investigate the relative abundance of metabolic and biogeochemical functional pathways in the prokaryotic community.

Metagenome reads were back-mapped to the catalogues using the BWA aligner [v0.7.17] [45] and unmapped reads were discarded. We considered a mapping quality threshold of 10. The number of aligned reads per gene was obtained using HTSeq [v0.13.5][46] (max-reads-in-buffer= 10^11^, stranded = no, nonunique = all). Counts per gene were normalized by gene length and the geometric mean abundance of 10 selected single-copy genes in each sample for prokaryotes [47] or metagenome size in gigabases for microbial eukaryotes. Normalized gene abundance tables were generated, including the abundance of each gene in each sample. The corresponding functional abundance tables were generated by adding all normalized abundances of all genes annotated to a specific KEGG function. Bioinformatics analyses were performed at the Marine Bioinformatics Service (MARBITS; https://marbits.icm.csic.es) of the Institute of Marine Sciences (ICM-CSIC) in Barcelona, Spain.

### Metagenome Assembled Genomes (MAGs)

Metagenomes were co-assembled using MEGAHIT [v1.2.8] [48]. Then, clean reads were mapped against the co-assembled contigs using BWA [v0.7.17] [45] to obtain contig abundance. Co-assemblies were binned using three binning programs: MetaBAT [v2.12.1] [v2.12.1] [55] (minContig = 2500), MaxBin [v2.2.5] [50] (min_contig_lenght = 2500, max_iteration = 50, prob_threshold = 0.9), and CONCOCT [v0.4.2] [51] (Length_threshold = 2500, clusters = 400) considering contigs with >2500 bp length. The three pools of bins were co-refined using Metawrap [v1.3-4bf5f8a] [52], considering genome completeness of >50% and contamination of <10% thresholds. Furthermore, COMEbin [v1.0.4] [53] was used as an additional binning algorithm with the default parameters. The constructed bins were dereplicated to select the best representative genomes, hereafter MAGs, using dRep [v2.3.2][54] with specific parameters: an Average Nucleotide Identity (ANI) threshold of 90% for the creation of primary clusters utilizing MASH and an ANI threshold of 99% for the formation of secondary clusters. Following these filtering steps, our dataset contained 15 high-quality prokaryotic MAGs. The taxonomic classification of MAGs was performed using GTDB-Tk [v1.5] [55]. Prokka [v1.14.6] [56] was used for functional annotation of MAGs. EnrichM [v0.5.0] [57] (DB = enrichm_database_v10, cutoff = 0.5) was used to investigate the KEGG functional modules’ completion in MAGs.

### Statistical analysis and visualizations

To analyze and visualize our data, we utilized the statistical computing environment ‘R’ (4.2.2). The data and tables were processed using the ‘Tidyverse’ [58]. The average abundance of the mTAG was calculated, and a 25th percentile threshold (P25) was applied to filter the data, retaining only values above this threshold for further analysis. The mTAGs data were normalized after filtering. The normalization process included calculating the sum of the counts of all filtered mTAGs across the samples. Subsequently, the counts of mTAGs were divided by the corresponding sum of all counts to obtain normalized values for further analysis. Relative abundance was calculated using the function ‘make_relative’ from the ‘ *Funrar*’ package [59]. Ecological statistical analyses were performed using the R packages ‘*Vegan*’ [60], and ‘*Ecodist*’ [61]. The Bray-Curtis dissimilarity was employed to perform non-metric multidimensional scaling (NMDS) of the samples according to their mTAG, gene, and KEGG profiles. Permutational multivariate analyses of variance (PERMANOVAs) were performed with 999 permutations to test for overall differences in the Bray-Curtis dissimilarity matrix between sample categories. Diversity was estimated using the Shannon index, and community richness was estimated using Chao1. The statistical significance between the two depth groups was determined using a *t*-test analysis. We used the ‘*MicroNiche*’ package [62] to assess the niche breadth. We analyzed and visualized the enrichment of KEGG pathways and modules using the R package ‘*clusterProfiler*’ [63]. We performed differential abundance analysis for the identified KEGG genes using ‘*lmFit*’ and ‘*eBayes*’ functions from the ‘*limma*’ package [64]. The results were visualized using ‘*ggplot2*’ [65] for statistical and heatmap plotting, and ‘*circlize*’ [66] for chord diagram plotting.

## Results

### Impact of depth on microbiome gene composition

In total, 430.6 Gb of sequencing data were obtained for the 10 *Eunicella singularis* metagenomes (approximately 3 × 10^8^ reads per sample). As expected, most of the reads were eukaryotic. The single-sample assemblies generated ca. 17.6x10^6^ contigs for all samples. We identified 12.3x10^6^ eukaryotic (70%) and 5.2x10^6^ prokaryotic contigs (30%). The number of eukaryotic and prokaryotic contigs was higher in the shallow samples. The average eukaryotic contig size in shallow samples was larger than that in mesophotic samples. In contrast, the mesophotic samples’ average prokaryotic contig size was larger than that in the shallow samples (Supplementary Figure 1; Supplementary Table 1).

After removing the predicted genes (ORFs) assigned to Anthozoa, the non-metric multidimensional scaling (NMDS) results of the eukaryotic genes (n= 266,741) indicated a significant impact of depth on gene composition (Figure 1A) (PERMANOVA, R^2^ = 0.6, *p* = 0.007). The shallow samples had a higher mean gene diversity (Shannon index = 12.0; SD = 0.03) than the mesophotic samples (10.7; SD = 0.04) [*t*-test *p*-value = 9.66 e-11] (Figure 1B). In addition, the shallow samples showed a higher mean Chao1 richness (196,679, SD = 2059) than the mesophotic samples (82,850, SD = 1894) [*t*-test *p*-value = 2.81e-13] (Figure 1C).

**Figure 1.**
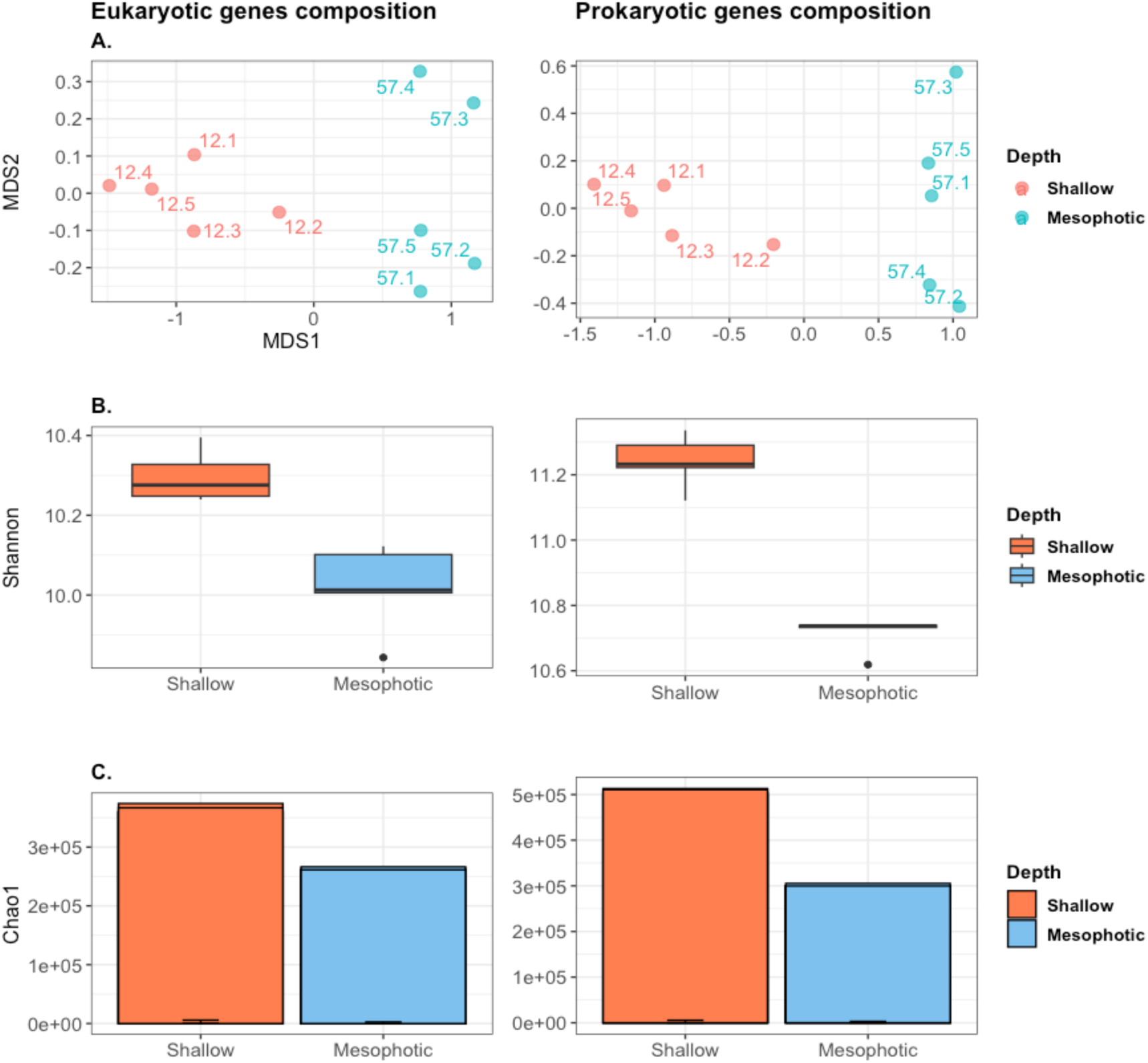
Gene (ORF) composition and diversity of the *Eunicella singularis* microbiome. **A.** Non-metric multidimensional scaling (NMDS) based on Bray-Curtis dissimilarities of eukaryotic and prokaryotic gene compositions. **B.** Box plot showing eukaryotic and prokaryotic gene diversity (Shannon index). **C.** Eukaryotic and prokaryotic gene richness estimates (Chao1).

Depth also influenced the prokaryotic gene (ORF) composition (PERMANOVA, R^2^ = 0.63, *p* = 0.009) (Figure 1A). The shallow samples had a higher average prokaryotic gene diversity (Shannon index, 13.0; SD = 0.03) than the mesophotic samples (12.3; SD = 0.04) [*t*-test *p*-value = 6.29e-06] (Figure 1B). The shallow samples had a higher average Chao1 gene richness (545,776.8, SD = 10554) compared to the mesophotic samples (332,615.6, SD = 7042) [*t*-test *p*-value = 2.6-e9] (Figure 1C).

### Taxonomic composition of the *Eunicella singularis* microbiome

The mTAG analysis based on the small subunit ribosomal RNA (SSU-rRNA) gene yielded 1,924,784 mTAGs. These reads consisted of 27% Opisthokonta (mostly gorgonian), 15% prokaryotes, 2% alveolates, and 1% other microeukaryotes, with the remaining 38% being unaligned or unassigned mTAGs. We excluded mTAGs identified as Cnidarian, unaligned, or unassigned from the downstream analysis. The Dinoflagellates’ mTAG-based NMDS results indicated clusters of samples corresponding to the two depths, indicating clear differences in Dinoflagellate composition between the shallow and mesophotic samples (PERMANOVA, median R^2^ = 0.78, *p* = 0.009) (Figure 2A). The shallow samples had higher dinoflagellate mTAGs richness and diversity (Chao1 μ = 249.4; Shannon μ= 3.30) than the mesophotic samples (Chao1 μ= 17.4; Shannon μ= 0.05) [Chao1 t-test *p* = 1.087e-05; Shannon t-test *p* = 0.0001] (Figure 2B, C). The high number of variants seen in shallow samples is most likely due to a combination of increased symbiont abundance and the multi-copy nature of the ribosomal marker, which may produce several sequence variants per cell.

**Figure 2.**
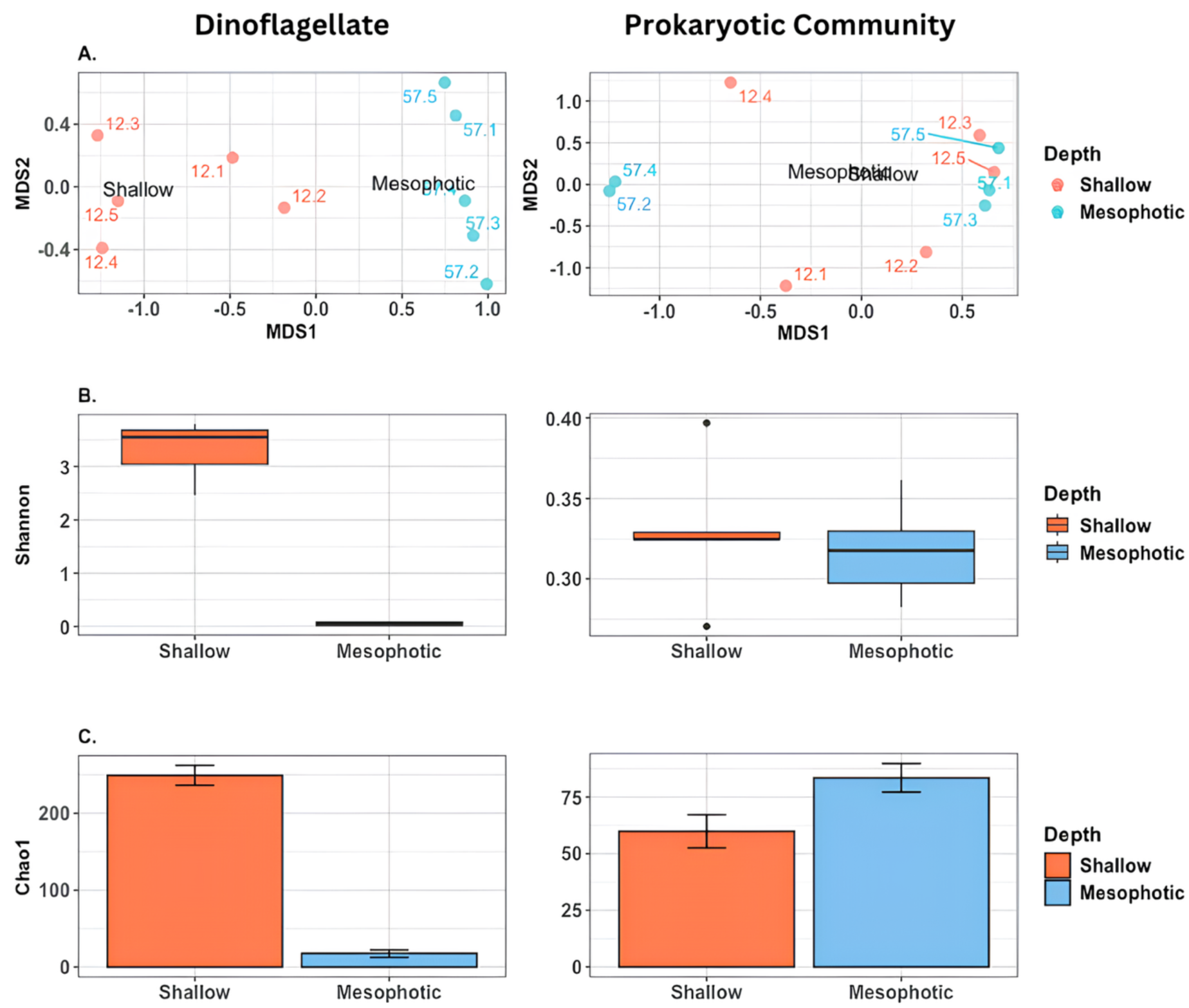
Composition and diversity of the *E. singularis* microbiome based on mTAGs. **A.** Non-metric multidimensional scaling (NMDS) based on Bray-Curtis dissimilarities (considering dinoflagellate [left panel] and prokaryotic [right panel] taxa). **B.** Diversity of shallow and mesophotic assemblages of dinoflagellates and prokaryotes (Shannon Index). **C.** Richness (Chao1) of shallow and mesophotic assemblages of dinoflagellates and prokaryotes.

The prokaryotic mTAG-based NMDS analysis did not reveal distinct clustering of samples according to depth, indicating no apparent differences in community composition between shallow and mesophotic samples (PERMANOVA, median R^2^ = 0.73, *p* = 0.09) (Figure 2A). Similarly, there was no significant difference in diversity between the two depths (shallow Shannon μ= 3.32; mesophotic Shannon μ= 0.31; *t*-test *p* = 0.65) (Figure 2B). In contrast, mesophotic samples exhibited significantly higher prokaryotic richness (Chao1 μ= 83.4) compared to shallow samples (Chao1 μ = 59.8; *t*-test *p* = 0.042) (Figure 2C).

The curated taxonomic annotation revealed that the gorgonian microbiomes comprised five main microbial phyla, including one dominant eukaryotic (Dinoflagellata) and several prokaryotic lineages. Within the protistan fraction, 33,279 mTAGs were assigned to Dinoflagellates (∼99% of all protistan mTAGs), with *Symbiodinium* as the dominant genus. After filtering out low-abundance reads (<50 total counts), the prokaryotic community was mainly represented by four bacterial classes: Alphaproteobacteria, Gammaproteobacteria, Bacilli, and Bacteroidia (Supplementary Table 2). *Endozoicomonas* dominated the bacterial communities at both depths, accounting for approximately 97% of the overall prokaryotic mTAGs. Despite the presence of different *Endozoicomonas* spp. mTAGs, only one taxon was consistently dominant across all samples (Supplementary Table 3). The genus *Bermanella* followed *Endozoicomonas*, comprising approximately 2% of the sequences. Other taxonomic groups, including the BD1-7 clade, *Sphingomonas*, *Mycoplasma*, *Thalassolituus*, *Halioglobus*, *Spiroplasma*, and *Aquimarina*, collectively constituted about 1% of the bacterial mTAGs (Figure 3, Supplementary Table 2).

**Figure 3.**
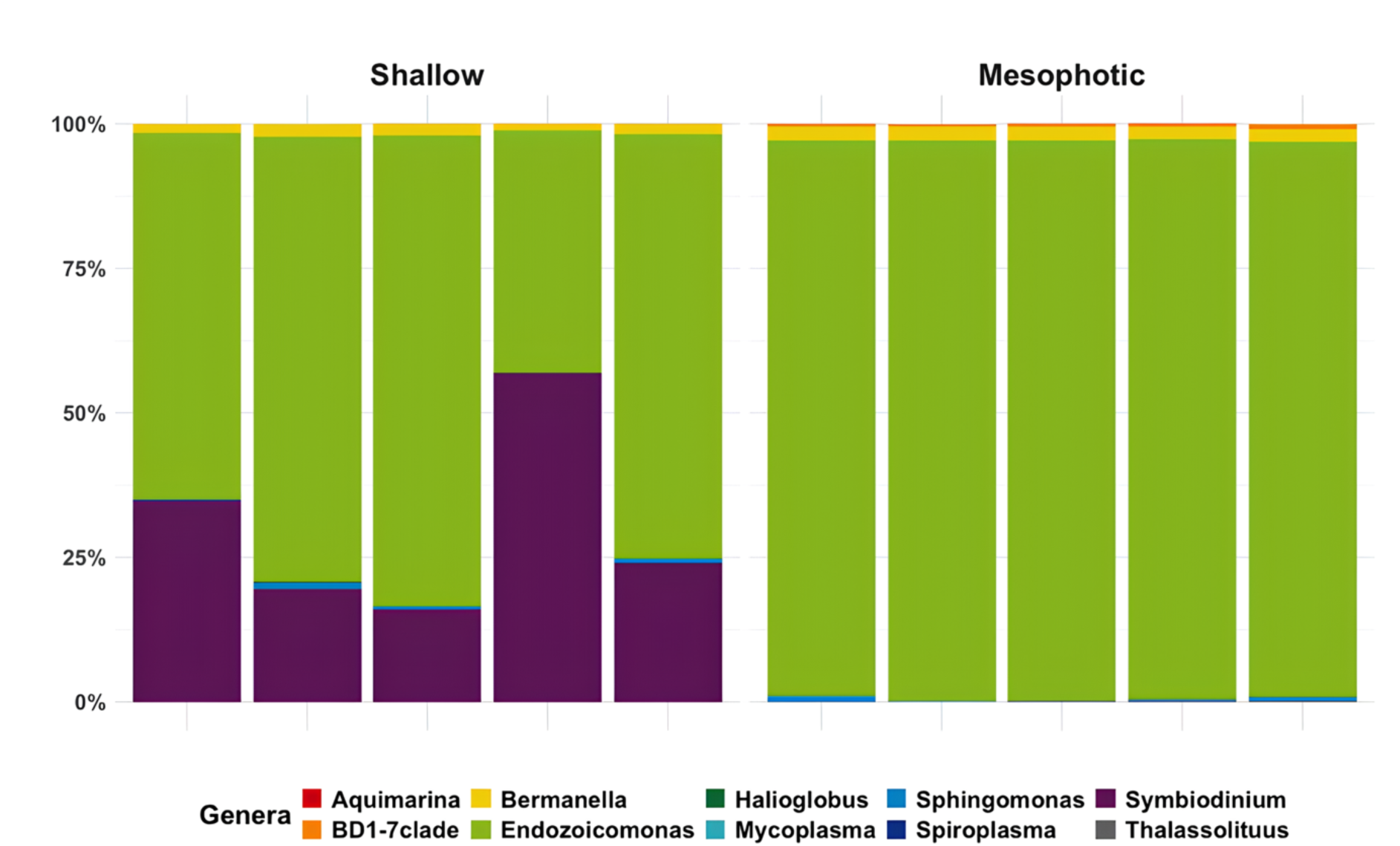
Taxonomic composition of the *E. singularis* microbiome based on mTAGs. The panels show the relative abundances of the most abundant genera in shallow (12m) and mesophotic (57m) samples.

We analyzed the niche breadth of microbial taxa across shallow vs. mesophotic samples using Levins’ approach [62]. Habitat generalist taxa were identified based on a broad distribution (Levins. Bn > 0.55) and no significant deviation from random expectation (*p* > 0.05). Conversely, taxa with narrower distributions (Bn ≤ 0.55) and significant deviation from random expectation (*p* < 0.05) were considered as habitat specialists. As a result, *Symbiodinium* and *Spiroplasma* were classified as shallow-water specialists, whereas BD1-7, *Thalassolituus*, and *Aquimarina* were identified as mesophotic specialists. The genera *Endozoicomonas*, *Bermanella*, *Halioglobus*, *Mycoplasma*, and *Sphingomonas* were identified as depth generalists, indicating that they were distributed across both depths (Supplementary Figure 2; Supplementary Table 4).

### Microbiome functional gene structure

Functional annotation of *Symbiodinium* genes resulted in 1,003 KEGG orthologs (KO’s) (hereafter KEGG). *Symbiodinium* functional genes were absent from the mesophotic samples in accordance with the results of the taxonomic analysis. We identified 53 metabolic pathways, most of which were shared with the prokaryotic community, except for three pathways: photosynthesis (ko00195), carbon fixation in photosynthetic organisms (ko00710), and nitrogen metabolism (ko00910) (Supplementary Table 5).

The functional annotation of prokaryotic genes resulted in 3,129 KEGG’s after filtering out low-abundance ones. Although no significant differences were observed in the overall composition between depths (PERMANOVA, p = 0.19), nor in alpha diversity metrics (Chao1 t-test p = 0.54; Shannon’s t-test p = 0.07) (Supplementary Figure 3), further analysis revealed notable patterns in functional enrichment. A total of 91 metabolic pathways were identified in the prokaryotic community, 82 of which were shared across depths. In addition, the prokaryotic community in mesophotic samples showed a higher abundance of KEGGs related mainly to four metabolic categories: carbohydrates, energy, amino acids, cofactors, and vitamin metabolism (Supplementary Figure 4). Some of the pathways that were found to have a higher abundance (p.adj < 0.05) in mesophotic samples were central metabolism (Glycolysis, Gluconeogenesis), oxidative phosphorylation (NADH-quinone oxidoreductase, cytochrome c oxidase (cbb3-type), and other prokaryotic cytochrome c oxidase), and the secretion system (Type III Secretion, Type II Secretion, Sec-SRP, and Twin arginine targeting) (Figure 4; Supplementary Table 6).

**Figure 4.**
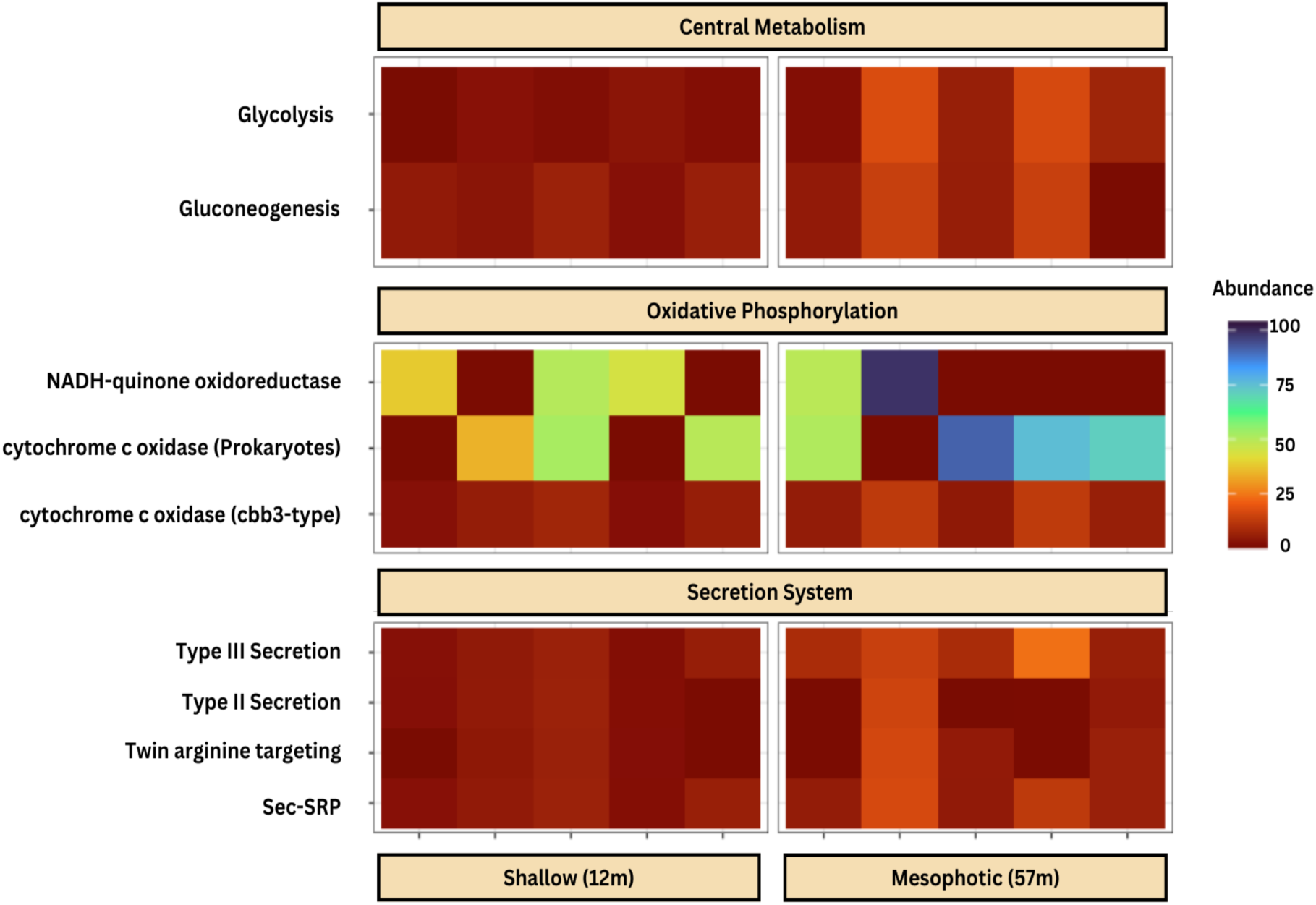
Relative abundances of specific prokaryotic metabolic pathways in the *Eunicella singularis* microbiome from shallow (12 m) and mesophotic (57 m) depths. Mesophotic samples showed higher oxidative phosphorylation pathways, particularly NADH–quinone oxidoreductase and cytochrome c oxidase, suggesting greater oxygen-dependent energy metabolism at depth. The type III secretion system pathway was also more abundant in mesophotic samples, indicating a possible interaction strategy with the coral host and surrounding microbiota. The used abundance unit (not shown) is the read counts per gene normalized by gene length and the geometric mean abundance of 10 selected single-copy genes in each sample.

Three KEGG pathways were found only in shallow samples, including photosynthesis (ko00195, ko00196) and phosphonate and phosphinate metabolism (ko00440) (q-value < 0.05) (Supplementary Table 5). Furthermore, six pathways were limited to the mesophotic prokaryotic community: steroid degradation (ko00984), pentose and glucuronate interconversion (ko00040), methane metabolism (ko00680), pinene, camphor, and geraniol degradation (ko00907), C5-Branched dibasic acid metabolism (ko00660), and glycerophospholipid metabolism (ko00564) (q-value < 0.05) (Supplementary Table 5).

We identified 49 KEGG modules, functional units within a metabolic pathway that execute specific biological functions [67], in the prokaryotic community, 44 of which were shared across different depths. However, four KEGG modules were enriched in shallow samples (q-value < 0.05). These modules include phylloquinone biosynthesis (M00932), NAD(P)H: quinone oxidoreductase (M00145), glycogen degradation - glycogen to glucose-6P (M00855), and lysine degradation - L-lysine to glutarate to succinate/acetyl-CoA (M00857). In contrast, only one module showed higher enrichment in mesophotic samples: catechol meta-cleavage (M00569).

We further aimed to identify potential metabolic pathways that were common but variable between prokaryotes and *Symbiodinium*. A total of 1,132 KOs were present in both *Symbiodinium* and prokaryotes. Among these KOs, 10 exhibited a noticeable abundance and varied significantly between the two depths (adj. P-value < 0.01). Some of these KOs were relatively more abundant in shallow samples. They are related to amino acid metabolism (K00558, K17398), carbohydrate and carbon metabolism (glycolysis/gluconeogenesis K01007, starch and sucrose metabolism K01179, and carbon fixation K01601(*rbcL*,*ccbL*)). Conversely, the mesophotic community showed higher relative abundances of K00232 (lipid metabolism), K00681 (amino acid metabolism), and K08678 (carbohydrate metabolism) (Figure 5; Supplementary Table 7).

**Figure 5.**
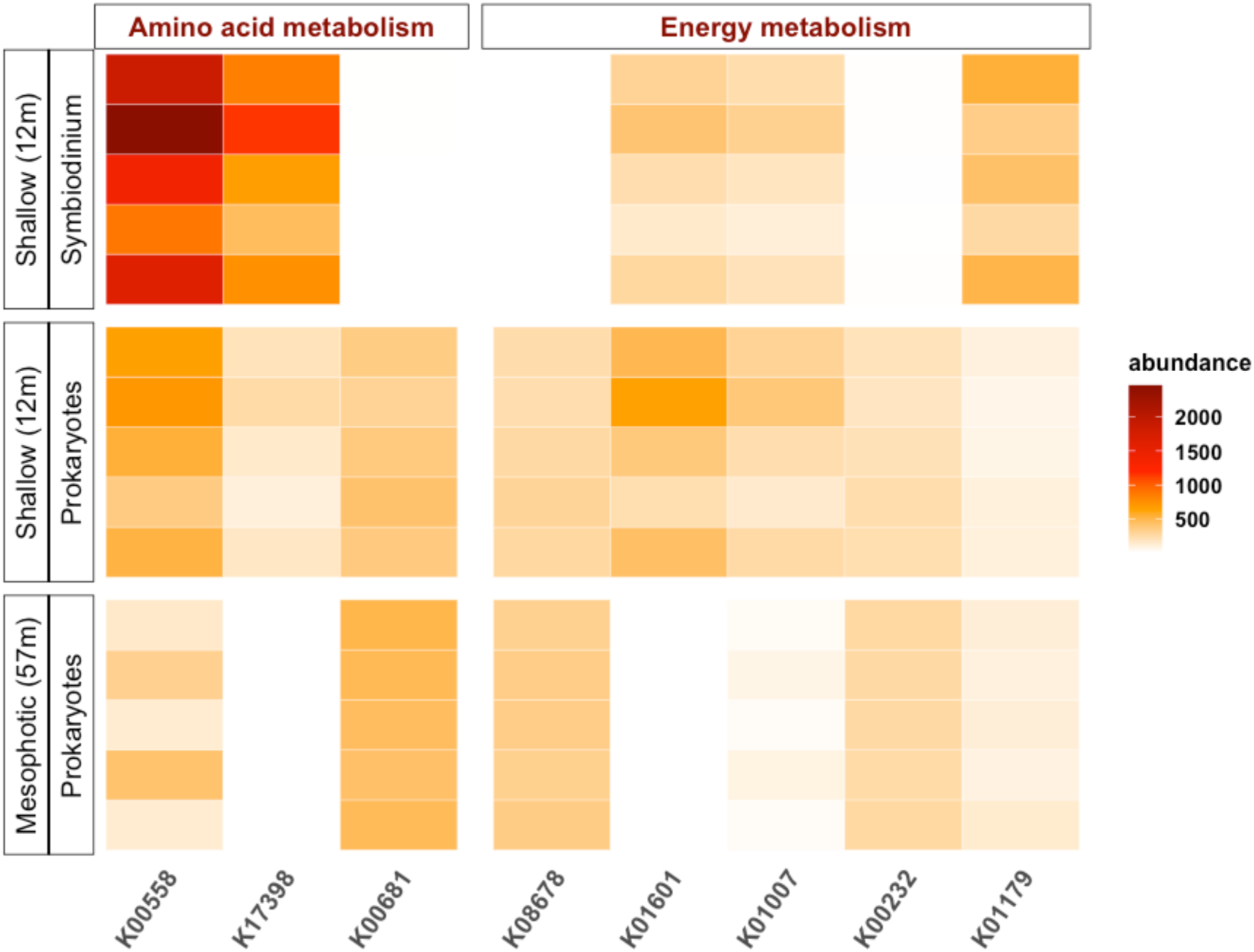
Abundances of KOs found in both *Symbiodinium* and prokaryotic communities of *Eunicella singularis* across depths. The abundance unit is the read counts per gene normalized by gene length and the geometric mean abundance of 10 selected single-copy genes in each sample.

In addition, we analyzed Open Reading Frames (ORFs) related to each of these functions. Interestingly, *Symbiodinium* showed a high number of ORFs annotated as DNA (cytosine-5)-methyltransferase 1 (K00558), DNA (cytosine-5)-methyltransferase 3A (K17398), and ribulose-bisphosphate carboxylase large chain (K01601) (Supplementary Figure 5). By contrast, the prokaryotic fraction showed relatively few ORFs annotated as K00558 and an almost complete absence of K17398 and K01601.

Finally, we compared the pathways shared between *Symbiodinium* and the associated prokaryotic community and found 52 shared pathways. The photosynthetic pathway (ko00195) was detected only in shallow samples, where it is most likely associated with *Symbiodinium*, indicating the expected absence of photosynthetic activity in the mesophotic community. In contrast, the prokaryotic community in the mesophotic samples showed higher KEGG diversity for the metabolism of cofactors and amino acids (Supplementary Figure 6).

### Metagenome Assembled Genomes (MAGs) analysis

We recovered 15 de-replicated prokaryotic MAGs with genome completeness values ranging between 66.28% and 99.65% (mean ∼ 83%) (Supplementary Table 8). These MAGs belonged to six bacterial classes: Cyanobacteria, GCA-001730085, Spirochaetia, Gammaproteobacteria, Alphaproteobacteria, and Bacilli. As expected, one MAG belonging to Endozoicomondaceae (MAG.25993) was dominant at all depths, being more abundant in the mesophotic. The other MAGs did not show statistical differences between the depths in terms of abundance (p < 0.05), except for MAG.12, Pseudomonadales (genus DT-91), which showed higher abundance in mesophotic samples despite its overall low abundance (Figure 6, Supplementary Table 8). Interestingly, one *Endozoicomonas* MAG (MAG.17) had a low abundance, which did not differ between depths and thus may not represent the dominant *Endozoicomonas* identified by mTAGs.

**Figure 6.**
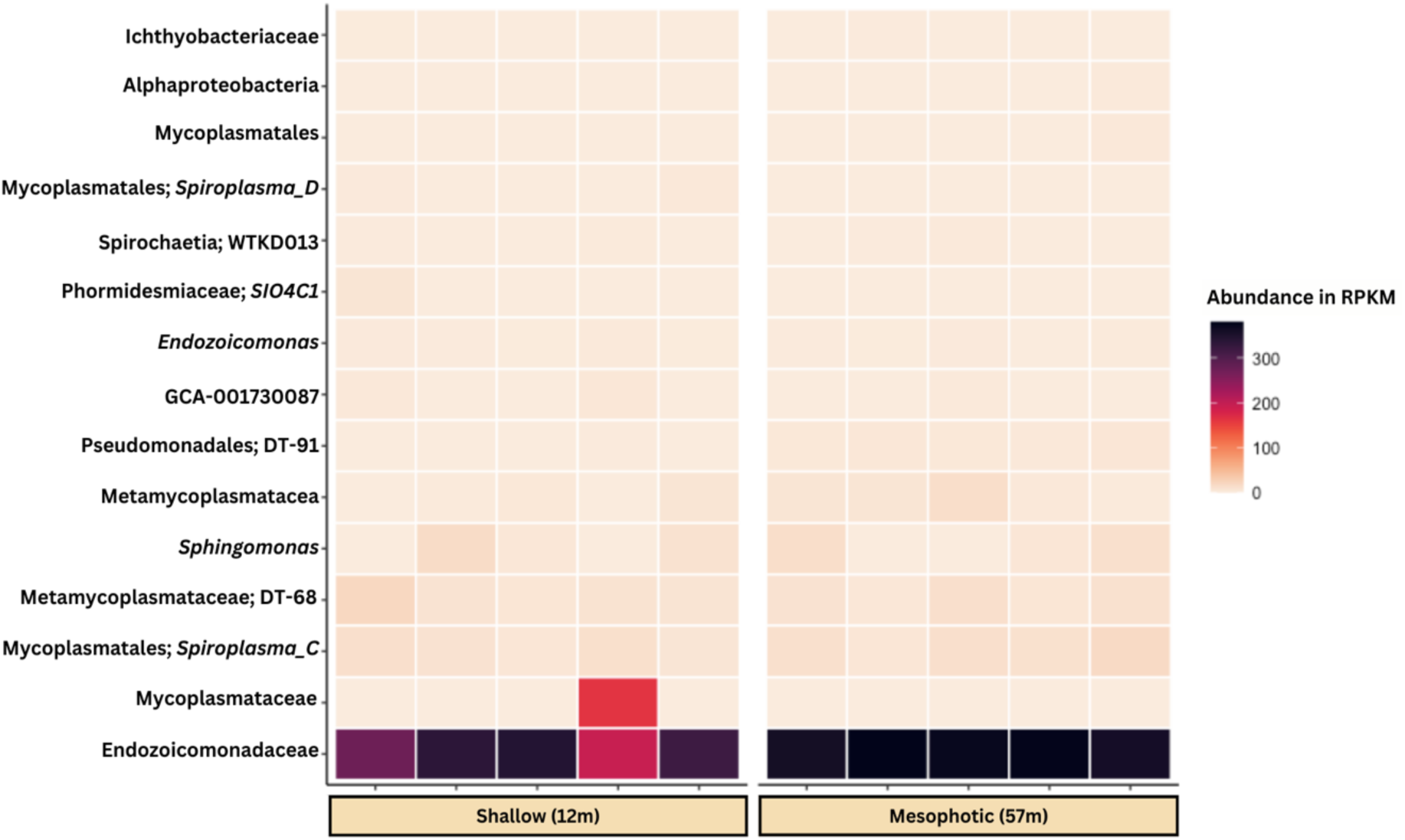
Abundance of the 15 prokaryotic MAGs. reconstructed from *E. singularis* metagenomes in shallow and mesophotic depths. RPKM = Reads Per Kilobase of MAG per Million of mapped reads

We further explored the potential contribution of MAGs to previously identified metabolic functions of the microbiome. While most MAGs encoded proteins involved in the central metabolism (Glycolysis, Gluconeogenesis), few MAGs encoded genes related to oxidative phosphorylation. For example, NADH-quinone oxidoreductase was found only in MAG.22 (*Sphingomonas*). In addition, the cytochrome cbb3-type was found in MAG.22 (*Sphingomonas*) and MAG.17 and MAG.25993 (Endozoicomondaceae). Cofactor and vitamin metabolism, such as menaquinone (K_2_), pantothenate (B_5_), pyridoxal (B_6_), biotin (B_7_), and cobalamin (B_12_), were found in only six MAGs: *Phormidesmiaceae* (genus SIO4C1), Pseudomonadales (genus DT-91), Pseudomonadales (genus *Endozoicomonas*), GCA-001730085, and Sphingomonadales (genus *Sphingomonas*). The MAG of the genus DT-91 (Order Pseudomonadales) had complete modules of vitamin and cofactor metabolism, such as biosynthesis of B_5_, B_6_, B_7_, and B_12_. In addition, Endozoicomondaceae MAGs exhibited biosynthesis modules for several vitamins, such as B_1_, B_6_, B_7_, and K_2_. Other MAGs also showed complete modules for vitamin and cofactor metabolism, including B_7_ and K_2_. Furthermore, eight MAGs were found to encode secretion system genes. For example, the twin-arginine translocation (Tat) system was found in eight MAGs, such as MAG.12 (Pseudomonadales, genus DT-91), MAG 17 and 25993 (Endozoicomondaceae), MAG.22 (Sphingomonas), MAG.220783 (Alphaproteobacteria), MAG.302726 (Ichthyobacteriaceae), MAG.309811 (GCA-001730086), and MAG.332111 (Phormidesmiaceae). Additionally, the type II secretion system was found in the MAG.12 (Pseudomonadales, genus DT-91), MAG 17 and 25993 (Endozoicomondaceae), MAG.22 (Sphingomonas). In contrast, the Type III secretion system was only found in MAG 17 and 25993 (Endozoicomondaceae) (Figure 7; Supplementary Table 9)

**Figure 7.**
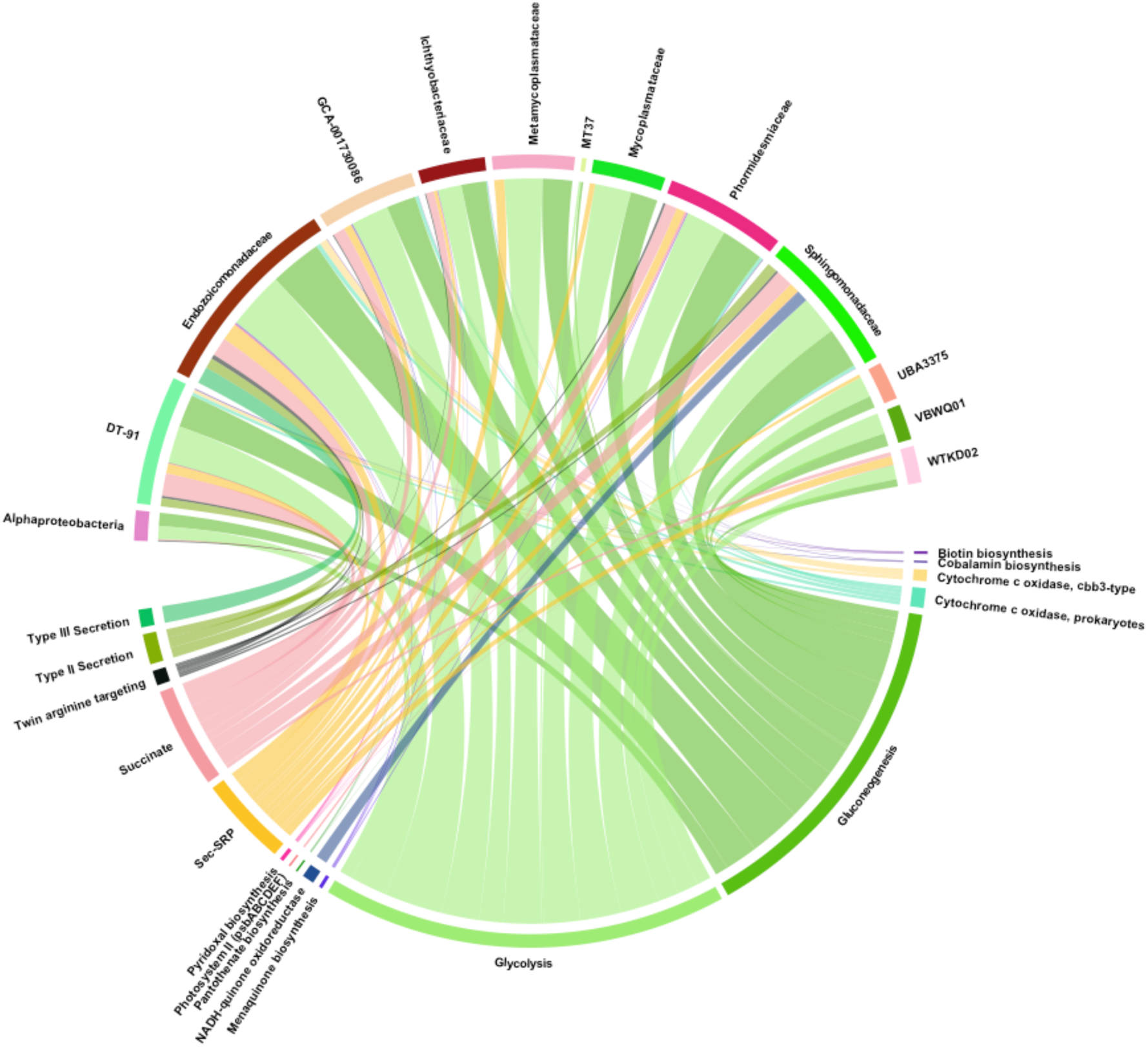
Chord graph for the 15 MAGs illustrating their contribution to metabolic pathways in the *E. singularis* microbiome. Each MAG is connected with pathways based on the KOs encoded in their genomes.

**Figure 8.**
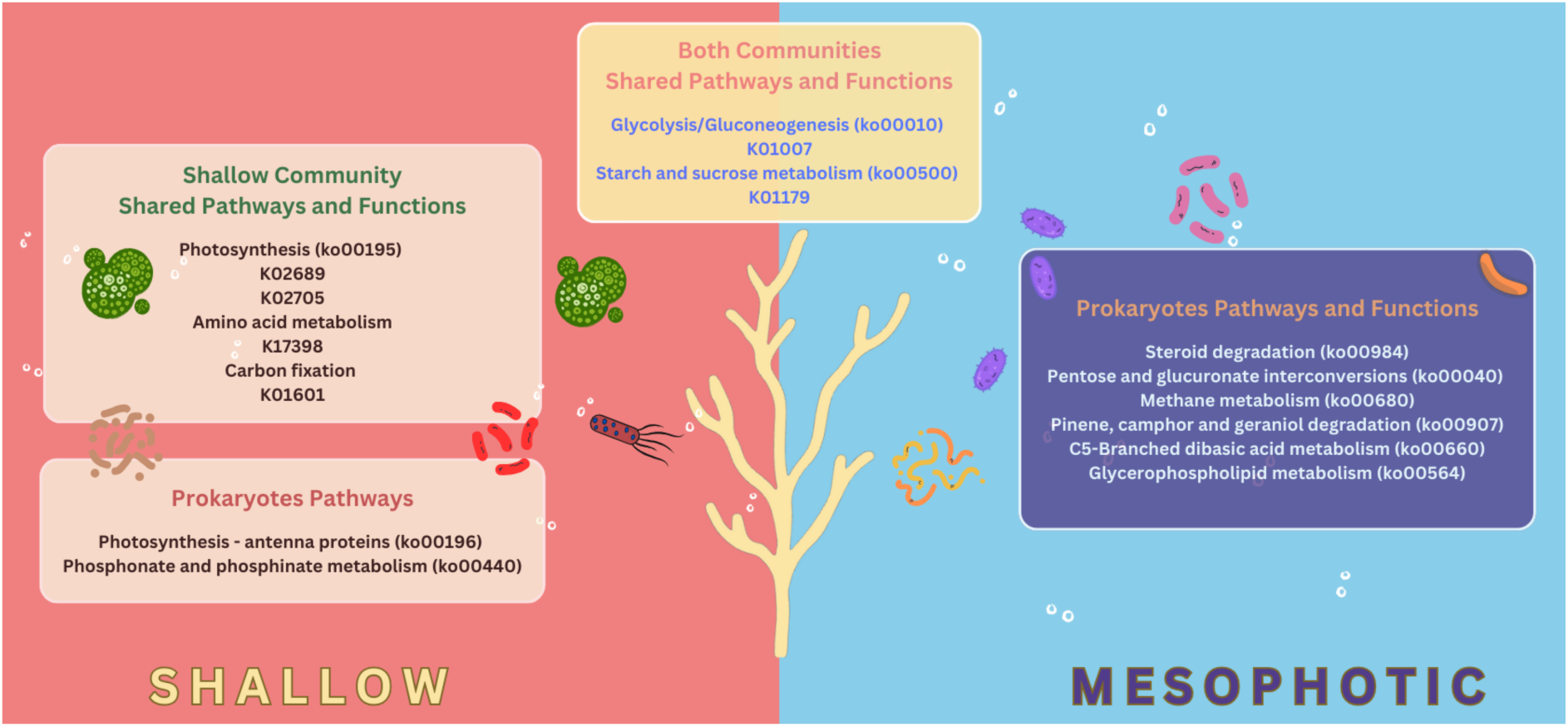
Summary of the unique metabolic features of shallow-water and mesophotic microbial communities associated with *E. singularis*. In shallow waters, the microbiome, particularly *Symbiodinium*, uses photosynthesis as a source of fixed carbon (left panel). In contrast, the mesophotic microbiome was enriched in specific metabolic pathways related to carbon metabolism, including steroid and aromatic hydrocarbon degradation, suggesting its adaptation to deeper, lower-light environments (right panel).

## Discussion

### The microbiome of Eunicella singularis changes in composition with depth

We found that E. singularis hosted fewer Symbiodinium in mesophotic samples and that its microbiome was richer and more diverse in shallow colonies. These differences are probably linked to the decline in light irradiance with depth. At mesophotic depths, photosynthetically active radiation (PAR) is approximately 25 times lower than at 15 m (1% vs. 25% of surface PAR, respectively [68]. In contrast, at the functional level, there was no significant difference on the composition of prokaryotic KEGG between shallow and mesophotic samples. Based on the mTAGs SSU-RNA taxonomic annotation, we found that *Symbiodinium* tended to dominate in the shallow *Eunicella singularis* colonies. Prokaryotic communities were prevalent in mesophotic metagenomes and were mainly dominated by Alphaproteobacteria and Gammaproteobacteria. Additionally, the Gamma-proteobacterial genera *Endozoicomonas* and *Bermanella* were the most abundant at both depths. *Endozoicomonas* is broadly distributed in marine environments across different depths and ecosystems [69], forming associations with various Cnidaria species [70], including Gorgonian corals [71]. Our findings suggest that the same *Endozoicomonas* mTAG dominates both shallow and mesophotic *Eunicella* microbiomes. This implies that the dominant bacterial species in the Mediterranean *E. singularis* remain constant along the depth gradient.

Previous studies have indicated that geographical distribution might impact the dominant Endozoicomonas OTUs in tropical gorgonian [14]. Further, four *Endozoicomonas* OTUs were reported previously to be associated with the temperate *E. singularis* across the Mediterranean at shallow depths [18]. It remains to be tested whether the dominant mTAG in this study matches the previously described OTUs or if they are specific to the Mediterranean region. Additionally, it is assumed that *Endozoicomonas*, in scleractinian corals, is involved in coral nutrient acquisition by participating in host-associated protein and carbohydrate transport and cycling, and structuring of the coral microbiome through the inhibition of pathogen proliferation using different mechanisms like the production of quorum-sensing antimicrobial compounds [71,72]. Despite their recognized mutualistic association with coral hosts, *Endozoicomonas* may adopt diverse roles, potentially residing within coral cells as commensals, parasites, or even pathogens [73].

Little is known about the role of *Bermanella* in the coral microbiome. One species belonging to this genus has been described, *Bermanella marisrubri*, which has been isolated from the Red Sea and described as a heterotrophic, marine, strictly aerobic, motile bacterium [74]. In addition, genomic evidence from *Bermanella* suggests a possible role in bioremediation and oil degradation by initiating *n*-alkane breakdown for other bacterial taxa, such as Spongiiabcteraceae, to carry out secondary *n*-alkane breakdown and beta-oxidization [75]. In addition, we identified other families of Alpha- and Gamma-proteobacteria frequently found in corals, including Saccharospirillaceae, Flavobacteriaceae, and Sphingomonadaceae [76]. We identified the Sphingomonadaceae genus *Sphingomonas* as a shallow specialist taxon in our dataset. Members of this family have been proposed to be putative coral pathogens [77].

Moreover, we identified rare (less abundant) taxa from two genera in the phylum Firmicutes: *Mycoplasma* and *Spiroplasma*. This phylum is assumed to include mutualistic or commensal microorganisms in temperate and deep-sea gorgonian species as well as in cold-water scleractinian corals [18,78,79]. It is also thought to be involved in the assimilation and dissimilation of sulfur reduction [80]. The genus *Mycoplasma* was observed in all samples and is commonly found in gorgonians [81,82] and cold-water scleractinians [83,84]. However, its precise functional role in corals and cnidarians remains unknown [85]. Interestingly, we identified *Spiroplasma,* known for its capacity to protect the host against pathogens [86], as a shallow-depth specialized taxon in our *Eunicella* samples. Some evidence showed that this taxon could confer protection against fungi through the production of unique toxins [86], suggesting its potential implication in the mechanisms of coral resilience against pathogens.

Specific bacterial taxa were strongly associated with depth. For instance, BD1-7 genus (family Spongiiabcteraceae) was more prevalent in mesophotic samples. This is consistent with its identification in the microbiomes of the mesophotic black corals *Antipathella subpinnata* and *Eunicella cavolini* at a depth of approximately 60 m [87]. It has also been recognized as a core member of the microbiome of *Eunicella singularis* [88]. In addition, Spongiiabcteraceae have been reported to be associated with the degradation of mono- and polycyclic aromatics [75].

Despite its low abundance, the genus *Thalassolituus* (family Saccharospirillaceae) was predominantly found in mesophotic colonies of *E. singularis*. It is assumed to be involved in the carbon and nitrogen cycles by utilizing acetate and C7-C20 aliphatic hydrocarbons for carbon, as well as ammonia and nitrate for nitrogen [89]. It has also been identified among associated bacteria in the endemic reef coral *Mussismilia braziliensis* [90]. However, we are unaware of previous studies that have addressed the presence of *Thalassolituus* in the *Eunicella singularis* microbiome. Similarly, *Aquimarina* (family Flavobacteriaceae) was identified as a mesophotic specialist taxon. Keller-Costa et al. [19] successfully isolated *Aquimarina* sp. strain EL33 from *Eunicella labiata* and obtained a nearly complete genome sequence. Analyses of the *Aquimarina* genome have revealed its potential roles in nutrient acquisition, utilization, and defense [19]. This finding offers valuable insights into the functional contribution of this bacterium to the holobiont. In addition to nutrient cycling, it has been suggested that it might inhibit pathogenic bacteria by producing exoenzymes, organic compounds, and bacteriocin [91].

Generally, the taxonomic analysis of the *E. singularis* microbiome revealed an absence of *Symbiodinium* at mesophotic depth, signaling a shift in microbiome composition. While the dominant bacterial species remained consistent, variation in the bacterial community was primarily observed among rare bacterial members (less abundant), with certain taxa showing depth specificity. In addition, the bacterial taxa identified as mesophotic depth specialists are thought to be capable of using different carbon sources, such as aromatic and hydrocarbon compounds [75].

### Changes of coral-microbiome functions with depth

As expected, the photosynthesis pathway (ko00195) was found only in the microbiome of shallow samples. Our mTAG results indicate that this is due to the absence of the endosymbiotic algae *Symbiodiniaceae* in the mesophotic samples, probably because of insufficient light irradiance at mesophotic depths[92]. We also found differences in abundance between depths for certain functional genes related to energy and amino acid metabolism. For instance, the ribulose-bisphosphate carboxylase large chain (RuBisCO large chain – *rbcL, cbbL*), which is related to carbon fixation in photosynthetic organisms, was prevalent in the shallow *E. singularis* microbiome as expected. Many *Symbiodinium* ORFs encoding RuBisCO were also present. This could be because *Symbiodinium* may have many gene copies encoding RuBisCO [93]. The prokaryotic community in the shallow *E. singularis* microbiome exhibited a high number of genes involved in carbon fixation. This suggests a relevant role of prokaryotes in providing fixed carbon to the coral. Until recently, only *Symbiodinium* was considered responsible for carbon fixation in coral holobionts [94]. This could also explain the presence of the autotrophic prokaryotic MAG *Phormidesmiaceae* (Cyanobacteria) in the microbiome.

Moreover, in the shallow samples, we found a high number of *Symbiodinium* ORFs that encode DNA methylation, particularly DNA (cytosine-5)-methyltransferase (K00558, K17398). DNA (cytosine-5)-methyltransferase is known to regulate gene expression and DNA maintenance [95]. One potential explanation for the abundance of these enzymes could be that dinoflagellates can hold multiple copies of a single gene [96], but also the nature of the endosymbiotic relationship between *Symbiodinium* and its hosts. *Symbiodinium* may adjust its gene expression in response to coral functions [97], but the specific mechanisms underlying this process remain unclear. Therefore, although DNA methylation likely contributes to these regulatory mechanisms, further research is needed to unravel the full extent of how *Symbiodinium* and gorgonians interact and influence each other’s gene expression and physiological processes.

Dinoflagellates exhibit a wide range of genome sizes up to 245×10^6^ kbp, containing 87,688 protein-coding and 92,013 total genes [98]. In the case of *Symbiodinium*, it has been reported to have ∼ 42,000 protein-coding genes and ∼19 exons per gene [99]. We have predicted a high number of *Symbiodiniaceae* genes (∼130k), which may be because we are predicting exons, and gene fragmentation could have inflated the number of *Symbiodiniaceae* genes.

We further explored how the prokaryotic microbiome might contribute to enhancing coral energy acquisition. The increased abundance of some pathways, such as cytochrome c oxidase, in the mesophotic community suggests higher energy production. In addition, the mesophotic community showed an enrichment of specific lipid metabolism functions, especially acyl-CoA oxidase. This allows fatty acids to be used as carbon sources [100] for energy production by the prokaryotic community. In addition, the enrichment of genes related to carbohydrate metabolism in the microbiome of the mesophotic *E. Singularis* colonies, such as UDP-glucuronate decarboxylase, could indicate higher biosynthesis of nucleotide sugars, such as UDP-xylose [101], which serves as a key precursor for the synthesis of diverse glycan structures and is essential for bacterial growth [102] and interaction with the host [103]. This may constitute a mechanism by which the prokaryotic community establishes colonization, avoids host immunity, or modulates the host response to pathogens [104].

Moreover, the mesophotic communities showed a higher abundance of genes related to amino acid metabolism, such as γ-glutamyltranspeptidase (GGT). This implies that the prokaryotic community uses GGT to degrade glutathione for amino acid utilization and as a nitrogen source. It can also be a virulence factor in some pathogenic gram-negative bacteria [105]. In addition, the enriched steroid degradation pathway in the mesophotic prokaryotic community suggests their ability to utilize organic compounds, such as steroids, as carbon sources. Furthermore, the presence of genes involved in the degradation of pinene, camphor, and geraniol within the *E. singularis* microbiome suggests that the prokaryotic community possesses the capability to utilize aromatic hydrocarbons as a carbon source [106], as well as the potential to degrade antibiotics. These observations suggest that the prokaryotic community associated with mesophotic *E. singularis* may have a distinct capacity to exploit diverse environmental carbon sources.

### MAG functional potential in the *E. singularis* microbiome

While glycolysis and gluconeogenesis were widespread among the *E. singularis* MAGs, oxidative phosphorylation genes were less common, with notable findings in specific taxa. For instance, NADH-quinone oxidoreductase genes have been exclusively identified in MAG.22 (*Sphingomonas*), highlighting its potential role in energy production. Moreover, the presence of cytochrome cbb3-type in MAG.22 (*Sphingomonas*), MAG.17, and MAG.25993 (*Endozoicomonadaceae*) suggest adaptations to varying oxygen levels [107].

Cofactor and vitamin metabolism genes were detected in a subset of MAGs, including Phormidesmiaceae, Pseudomonadales, and Sphingomonadales, indicating their importance in microbial growth and metabolism. Vitamin B plays a central metabolic role in marine bacteria and is important for nutrient uptake [108]. We detected a variety of vitamin B biosynthesis pathways in MAGs, including pantothenate (B_5_), pyridoxal (B_6_), biotin (B_7_), and cobalamin (B_12_). The recovery of a complete B_12_ biosynthesis module in MAGs from the genus DT-91 (Order Pseudomonadales) and a nearly complete module (85%) in SIO4C1 (Order Phormidesmiaceae) strongly suggests that these taxa can contribute to cobalamin provisioning within the coral holobiont. The B_12_ is an important vitamin for the coral and its symbiont *Symbiodinium*, as both lack cobalamin biosynthesis capability [109], which is required for methionine synthesis in the coral hosts [94]. This implies a potential role of both Pseudomonadales and Phormidesmiaceae in supporting corals with B_12_.

Moreover, some taxa showed potential for the biosynthesis of pantothenate (B_5_), such as Endozoicomonadaceae MAGs and GCA-001730085. Vitamin B_5_ has been suggested to play a role in energy generation as a precursor of coenzyme A, in addition to its use by commensal bacteria to enhance the protective activity against pathogens [110]. However, its precise role in the coral microbiome remains unknown. In addition, several MAGs had the potential to contribute to B_7_ biosynthesis, which is essential for bacterial metabolism, particularly in processes such as gluconeogenesis and fatty acid synthesis [111].

Additionally, the presence of type II and III secretion systems in three MAGs (DT-91, *Endozoicomonas*, and *Sphingomonas*) indicated their possible role in pathogenesis. The type II secretion system is a multiprotein system that translocates substances through the outer bacterial membrane, including virulence factors, enzymes, and effectors [112]. TSS III functions as an injectisome, allowing these bacteria to manipulate targeted cells and facilitate their transport across their membranes [113]. *Sphingomonas* was previously identified as a putative coral pathogen, which may explain the presence of these virulence factors in the genome. However, *Endozicomonas* is recognized as a coral symbiont, but lately, the nature of this relationship has been questioned [73]. One possible explanation is that *Endoziocomonas* may have an “accidental” pathogenic mode. To illustrate, the T3SS present in accidental pathogens serves as a fitness element in benign associations with primary hosts and as a virulence factor in accidental hosts. These pathogens may reside harmlessly in the hosts but can cause damage when encountering accidental hosts, although infection typically does not facilitate pathogen spread [114]. This could also explain the low abundance of these pathways compared to the dominance of *Endoziocomonas* in the microbiome.

## Conclusion

*Eunicella Singularis*, a soft gorgonian coral species commonly found in the Mediterranean Sea, possesses a microbiome that shows changes in taxonomic composition and function with depth. In shallow coral samples, the microbial community was predominantly composed of Symbiodiniaceae, which explains the prevalence of photosynthesis and carbon fixation genes. In addition, specific bacterial taxa, such as Spiroplasma, showed shallow depth preference. In contrast, the mesophotic Eunicella samples showed a higher prevalence of prokaryotes, which could explain the presence of more bacterial taxa with mesophotic depth niches, such as Thalassolituus, the BD1-7 clade, and Aquimarina. Moreover, the mesophotic microbiome showed a higher abundance of metabolic pathways related to carbon metabolism, suggesting its ability to adapt to a variety of alternative sources of carbon. In addition, they showed a higher abundance of genes related to cofactors and energy metabolism, suggesting higher microbial activity. Altogether, these patterns indicate that the transition from algal to microbial partners across depth gradients reflects a functional shift in the *E. singularis* holobiont—from phototrophic to heterotrophic support systems—enhancing its ecological plasticity and resilience.

## Acknowledgments

We would like to express our deep appreciation to King Abdulaziz University, Jeddah, Saudi Arabia, and the Saudi Cultural Mission in Madrid, Spain, for their support, which enabled us to conduct this research. Furthermore, we would also like to acknowledge the assistance provided by the Marine Bioinformatics Service (MARBITS; https://marbits.icm.csic.es) at the Institute of Marine Sciences (ICM-CSIC) in Barcelona, Spain. Finally, we extend our special thanks to the ECOgeochemistry of Benthic Environments Laboratory at Sorbonne University in Banyuls-sur-Mer, France, for their valuable contributions and collaboration in this scientific endeavor. A special thanks to Bruno Hesse and Jean Claude Roca for the support in deep samples collection.

## Conflict of interest

The authors declare no conflict of interest.

## Data availability

All data generated or analysed in this study are included in the manuscript

## Notes

### Competing Interest Statement

The authors have declared no competing interest.

