## Supplementry Figures for "Microbiome composition and function vary with depth in the Mediterranean gorgonian *Eunicella singularis*"

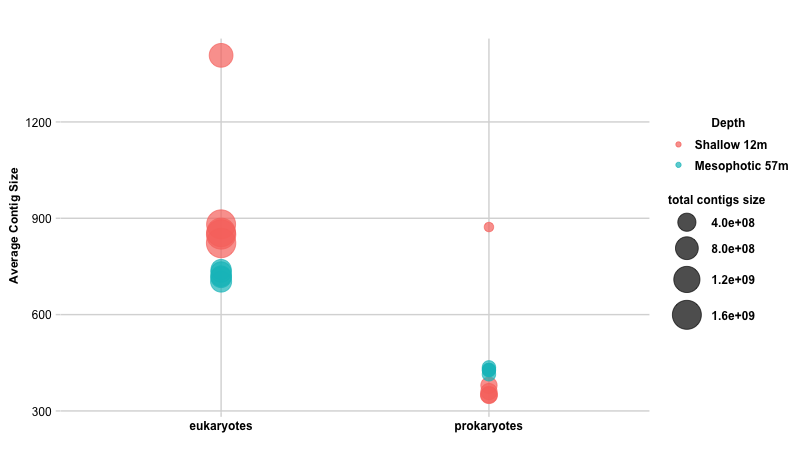

**Supplementary Figure 1**. Comparison of eukaryotic and prokaryotic contig sizes in shallow and mesophotic coral samples. Shallow metagenomes featured larger eukaryotic contigs, while mesophotic metagenomes exhibited slightly larger prokaryotic contigs compared to those in shallow samples.

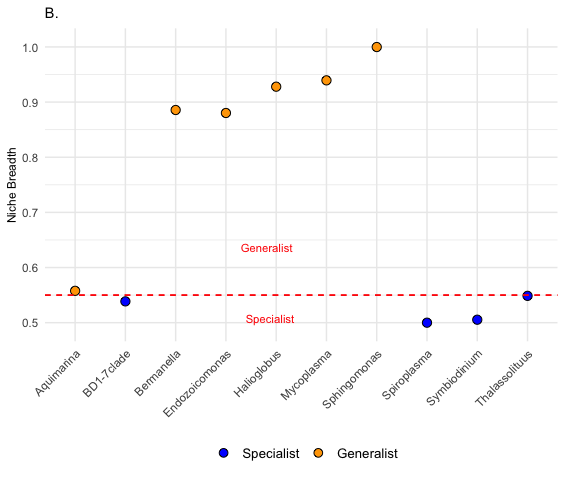

**Supplementary** **Figure 2.** Niche Breadth of the most abundant microbial taxa across shallow vs. mesophotic samples using Levins’ (Bn) approach. Taxa with Levins. Bn > 0.55 were identified as generalists while taxa with Levins. Bn ≤ 0.55 were identified as specialists.

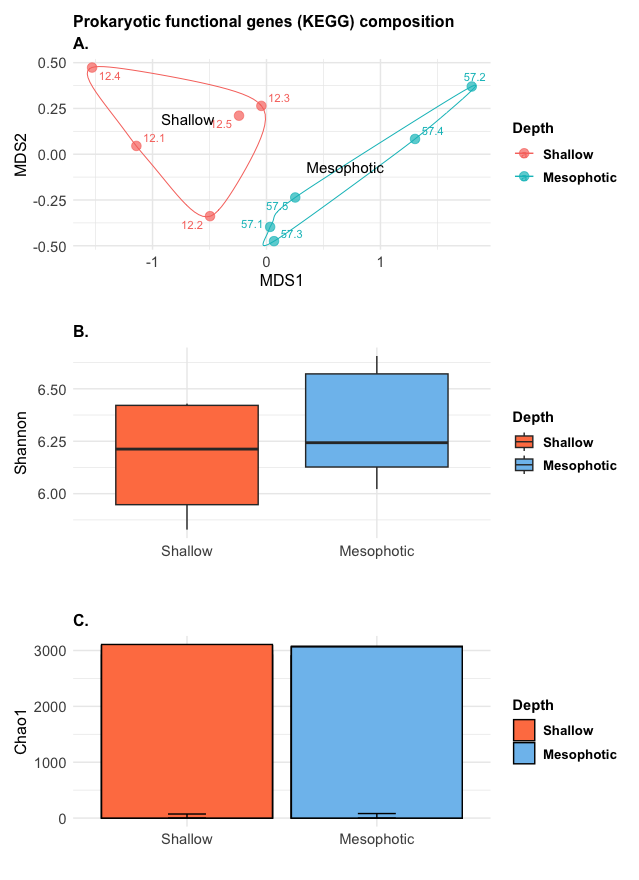

**Supplementary** **Figure 3.** Prokaryotic KEGG functions at both depths. The ordination shows variation in the prokaryotic KO composition between depths based on Bray Curtis distance (Panel A). However, no differences between surface and mesophotic samples were found in the Shannon index (Panel B) and Chao1 (Panel C).

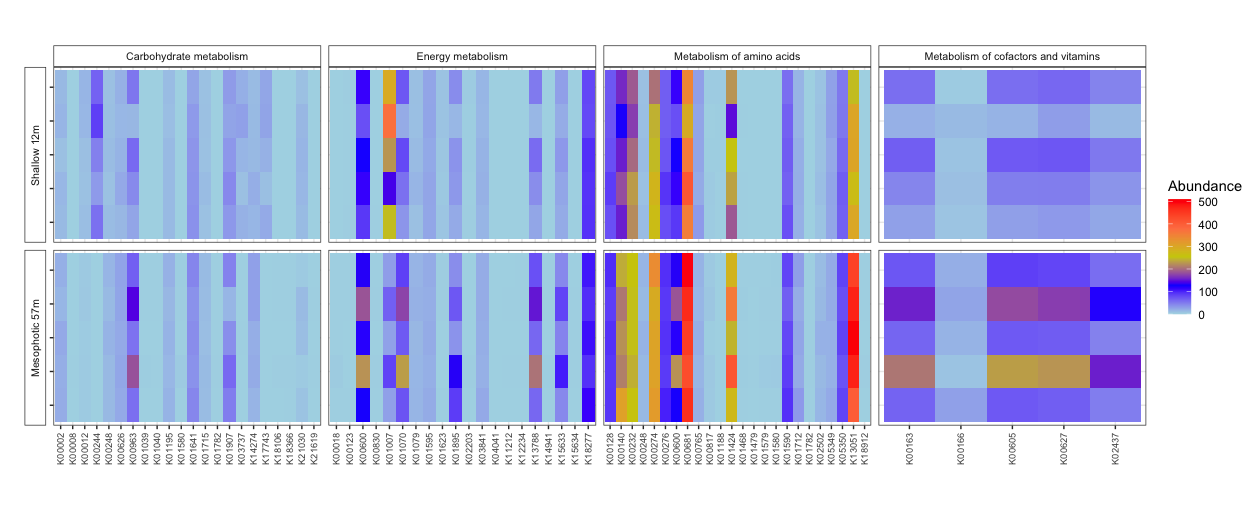

**Supplementary** **Figure 4.** Enrichment analysis of prokaryotic KEGG pathways in four metabolic categories: carbohydrate metabolism, energy metabolism, amino acid metabolism, and cofactor and vitamin metabolism between the shallow and mesophotic samples.

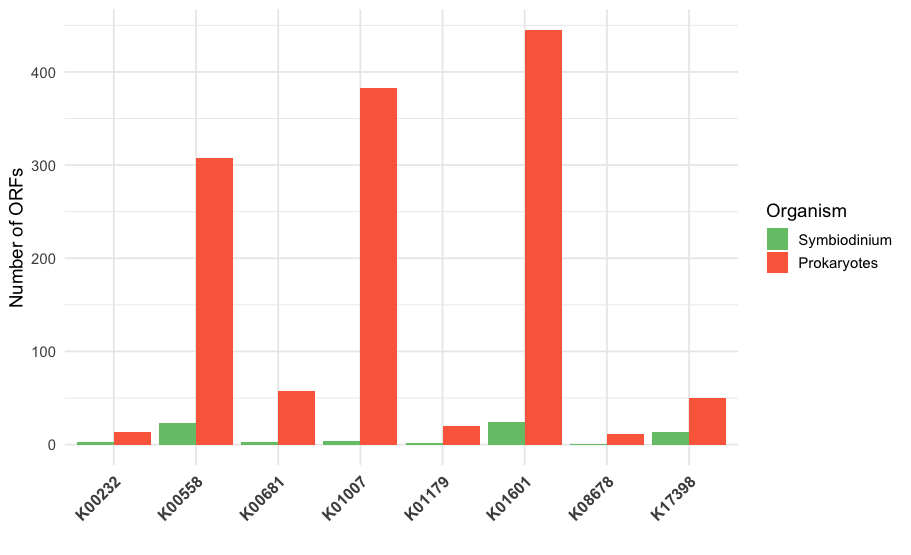

**Supplementary** **Figure 5.** The number of genes (ORFs) annotated for each shared KO in *Symbiodinium* and prokaryotic communities.

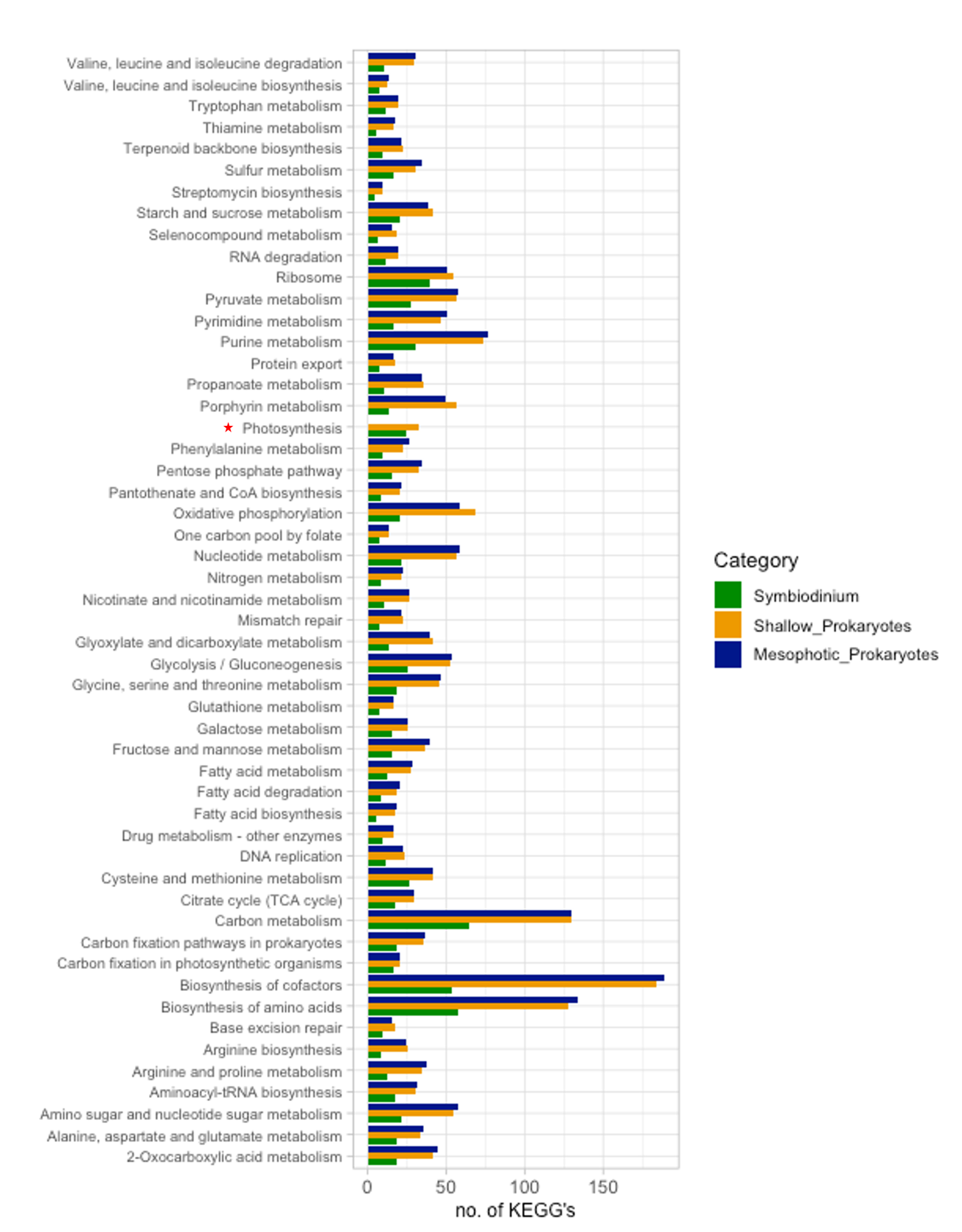

### **Supplementary** **Figure 6.** Shared enriched KEGG pathways between *Symbiodinium* and the prokaryotic community in the *Eunicella* microbiome. The photosynthesis pathway (marked with a star) was only found in shallow communities. However, the mesophotic microbiome showed more KEGGs related to the metabolism of cofactors and vitamins, and amino acid biosynthesis.
